# Cell-free DNA Profiling Informs Major Complications of Hematopoietic Cell Transplantation

**DOI:** 10.1101/2020.04.25.061580

**Authors:** Alexandre Pellan Cheng, Matthew Pellan Cheng, Conor James Loy, Joan Sesing Lenz, Kaiwen Chen, Sami Smalling, Philip Burnham, Kaitlyn Marie Timblin, José Luis Orejas, Emily Silverman, Paz Polak, Francisco M. Marty, Jerome Ritz, Iwijn De Vlaminck

## Abstract

Allogeneic hematopoietic cell transplantation (HCT) provides effective treatment for hematologic malignancies and immune disorders. Monitoring of post-transplant complications is critical, yet current diagnostic options are limited. Here, we show that cell-free DNA (cfDNA) in blood is a highly versatile analyte for monitoring of the most important complications that occur after HCT: graft-versus-host disease (GVHD), a frequent immune complication of HCT; infection; relapse of underlying disease; and graft failure. We demonstrate that these different therapeutic complications can be informed from a single assay, low-coverage bisulfite sequencing of cfDNA, followed by disease-specific bioinformatic analyses. To inform GVHD, we profile cfDNA methylation marks to trace the cfDNA tissues-of-origin and to quantify tissue-specific injury. To inform on infections, we implement metagenomic cfDNA profiling. To inform cancer relapse, we implement analyses of tumor-specific genomic aberrations. Finally, to detect graft failure we quantify the proportion of donor and recipient specific cfDNA. We applied this assay to 170 plasma samples collected from 27 HCT recipients at predetermined time points before and after allogeneic HCT. We found that the abundance of solid-organ derived cfDNA in the blood at one-month after HCT is an early predictor of acute graft-versus-host disease (area under the curve, 0.88). Metagenomic profiling of cfDNA revealed the frequent occurrence of viral reactivation in this patient population. The fraction of donor specific cfDNA was indicative of cell chimerism, relapse and remission, and the fraction of tumor specific cfDNA was informative of cancer relapse. This proof-of-principle study shows that cfDNA has the potential to improve the care of allogeneic HCT recipients by enabling earlier detection and better prediction of the complex array of complications that occur after HCT.

## INTRODUCTION

More than 30,000 patients undergo allogeneic hematopoietic cell transplants (HCT) worldwide each year for treatment of malignant and nonmalignant hematologic diseases^1–3^. Yet, myriad complications occur in this patient population. For example, up to 50% of patients experience graft-versus-host disease (GVHD), an immune response in which donor immune cells attack recipient tissues in the first year after transplantation^2,4–6^. Complications due to infection also occur frequently, mostly in the first year after transplantation, with bacterial and viral infection occurring in 52% and 57.9% of patients, respectively^7,8^. In addition, up to 50% of patients treated for malignant hematologic diseases suffer cancer relapse^9,10^. Last, graft failure is a major complication of HCT^11–14^.

Patient monitoring for post-HCT complications relies on a complex combination of diagnostic assays. Early and accurate diagnosis of GVHD is critical to inform treatment decisions and to prevent serious long-term complications. Unfortunately, there are few, noninvasive diagnostic options that reliably identify patients early after the onset of GVHD symptoms: In current practice, diagnosis of GVHD relies primarily on clinical symptoms and requires confirmation with invasive procedures, such as biopsies of the gastrointestinal tract, skin, or liver^15^. Furthermore, there is a critical need for tools that can broadly and sensitively inform infection. A wide range of microorganisms can cause disease after HCT, and infection testing currently relies on a combination of bacterial culture that are slow and suffer from a high false negative rate, and viral PCRs which have limited multiplexity. To screen for cancer recurrence, the presence of cancer cells in the circulation is used as a prognostic marker for relapse and disease-free survival. Current monitoring options of minimal residual disease include flow-cytometry, and quantitative PCR. However, these technologies are insensitive to genetic and phenotypic changes^16^. Donor chimerism is currently used to quantify engraftment, relapse and graft loss, but relies on the analysis of living cells, and may not be sensitive to the high turnover rate of leukemic cells^17^.

Here, we investigate the utility of circulating cell-free DNA as a versatile analyte to monitor HCT recipients after transplantation. Cell-free DNA in the blood of HCT recipients is a complex mixture of DNA from several sources: different tissues, microbes, donor cells and tumor cells^18–20^ (**Fig. 1A**). In this work, we demonstrate that a single assay, genome-wide methylation profiling of cell-free DNA, enables simultaneous monitoring of the major complications that arise after HCT. First, we show that methylation profiling by whole-genome bisulfite sequencing of cfDNA can be used to quantify the tissues-of-origin of cfDNA to thereby detect and quantify tissue injury due to GVHD after HCT. Second, we demonstrate the possibility to identify infectious agents via whole-genome bisulfite sequencing of cfDNA. Last, we show that the levels of donor- and tumor-derived cfDNA can inform engraftment, mixed chimerism, and cancer relapse. Together, this study provides a proof of principle that cfDNA profiling can be used to simultaneously monitor immune, cancer and infectious complications and treatment failure after allogeneic HCT.

**Figure 1.**
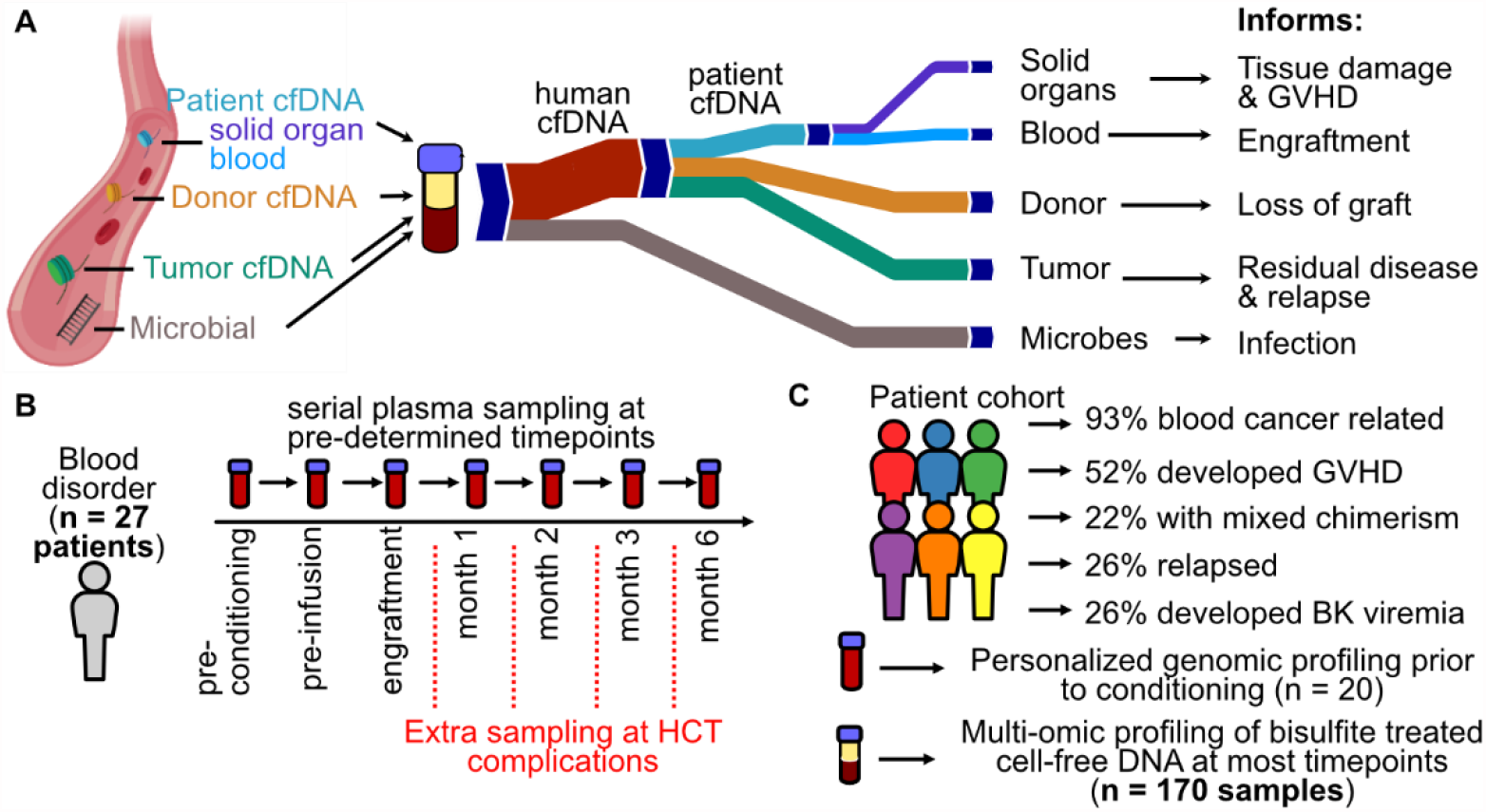
Study overview. **A** Cell-free DNA origins inform diverse transplant events and complications. **B** Plasma from 27 HCT recipients was serially collected at 7 or more predetermined timepoints. **C** Patient cohort characteristics.

## RESULTS

We performed a prospective cohort study to evaluate the utility of cfDNA to predict and monitor GVHD, infection, cancer relapse and treatment failure after allogeneic HCT. We selected 27 adults that underwent allogeneic HCT and assayed a total of 170 serial plasma samples collected at seven predetermined time points, including before conditioning chemotherapy, on the day of but before hematopoietic cell infusion, after neutrophil engraftment (>500 neutrophils per microliter), and at one, two, three and six months post HCT (**Fig. 1B**). Additional samples were collected at the time of presentation of complications, such as symptoms of BK disease. The test cohort included patients with both malignant (n=25) and non-malignant blood disorders (n=2) (**Fig. 1C, supplementary table 1**). Prior to conditioning, patient tumor cells were genotyped using a targeted deep sequencing panel^21^ (n = 20, total, 6 patients with copy number alterations). In total, 14 patients developed acute GVHD (GVHD+) and 13 did not (GVHD-), four patients experienced graft failure, seven developed BK virus viremia, and five patients suffered cancer recurrence (**Fig. 1C**, see Methods and SI).

We isolated cfDNA from plasma (0.5 mL-1.9 mL per sample) and implemented whole-genome bisulfite sequencing and bioinformatic analyses to profile epigenetic and genetic marks within cfDNA that may inform the diverse complications that arise after HCT. We implemented a single-stranded DNA (ssDNA) library preparation to obtain sequence information after bisulfite conversion^22,23^. This ssDNA library preparation avoids degradation of adapter-bound molecules which is common for WGBS library preparations that rely on ligation of methylated adapters before bisulfite conversion and avoids amplification biases inherent to WGBS library preparations that implement random priming^24^. We obtained 39 ± 14 million paired-end reads per sample, corresponding to 0.96 ± 0.4 fold per-base human genome coverage and achieved a high bisulfite conversion efficiency (99.4% ± 0.4%). We used paired-end read mapping to characterize the length of bisulfite treated cfDNA at single-nucleotide resolution and to investigate potential degradation of cfDNA due to bisulfite treatment. This analysis revealed a fragmentation profile similar to the fragmentation profile previously reported for plasma cfDNA that was not subjected to bisulfite treatment^25^. The mode of fragments longer than 100bp was 165 bp ± 7 bp, and Fourier analysis revealed a 10.4 bp periodicity in the fragment length profile (**supplementary figure 1)**. A second peak at 60-90 bp in the fragment length profile is characteristic of single-stranded library preparation methods and was reported previously^22,26^. Overall, we do not find evidence of significant cfDNA fragmentation due to bisulfite treatment, in line with a recent report^27^.

### Temporal dynamics of cell-free DNA tissues-of-origin in response to conditioning therapy and HCT

We first examined the utility of cfDNA tissues-of-origin by methylation profiling to identify organ injury due to GVHD after HCT in plasma samples obtained prior to the clinical diagnosis of GVHD. To quantify the relative proportion of cfDNA derived from different vascularized tissues and hematologic cell types we analyzed cfDNA methylation profiles against a reference set of methylation profiles of pure cell and tissue types^28–32^ (samples with sequencing depth greater than 0.1x, 138 reference tissues, see Methods, **Fig. 2A**,**B** and **supplementary dataset 1**). We computed the absolute concentration of tissue-specific cfDNA by multiplying the proportion of tissue-specific cfDNA with the concentration of total host-derived cfDNA (Methods). The most striking features seen in the data include: ***i***) a decrease in blood-cell specific cfDNA in response to conditioning therapy performed to deplete the patient’s hematopoietic cells (**Fig. 2C**), ***ii***) an increase in total cfDNA concentration at engraftment (**Fig. 2D**), ***iii***) a decrease in total cfDNA concentration after 180 days **(supplementary figure 2)**), and ***iv***) an association between tissue-specific cfDNA and the incidence of GVHD (see statistical analysis below).

**Figure 2.**
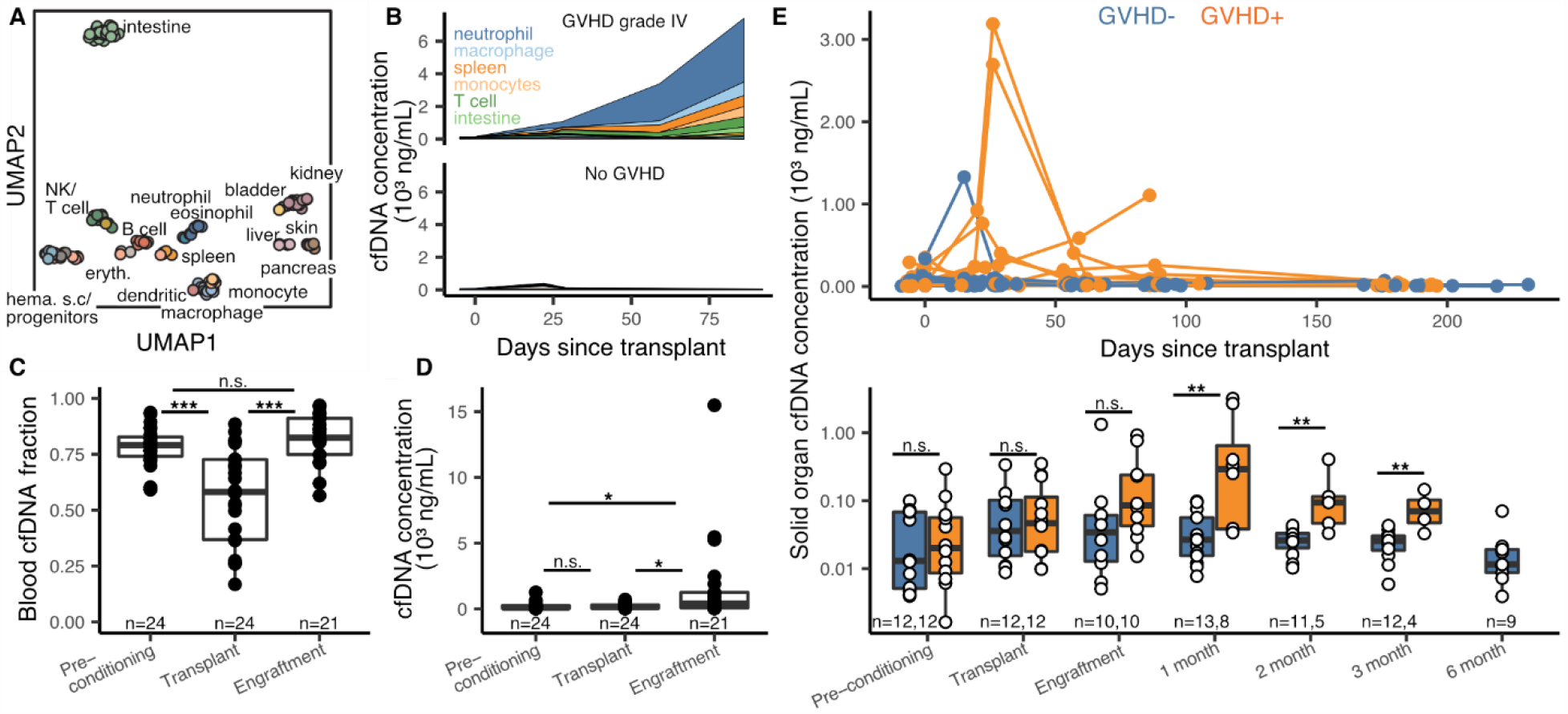
Host-derived cell-free DNA dynamics before and after HCT. **A** UMAP dimensional reduction of cell and tissue methylation profiles. Individual tissues are colored by UMAP coordinates using a linear gradient where each of the four corners is either cyan, magenta, yellow or black. **B** Examples of cfDNA dynamics in the case of severe GVHD (patient 003, top) and no GVHD (patient 017, bottom) in the first 3 months post-transplant. **C, D** Effect of conditioning and HCT infusion on cfDNA composition (C) and absolute concentration (D). **E** Solid-organ derived cfDNA concentration in plasma. Top: solid-organ cfDNA and days post-transplant for each patient time point. Bottom: solid organ cfDNA by time point. Samples are removed from analysis if plasma was collected after GVHD diagnosis. * p-value < 0.05; ** p-value < 0.01; *** p-value < 0.001

### Cell-free DNA tissues-of-origin by methylation profiling to monitor GVHD

We next examined these features in more detail to explore the utility of these measurements to monitor immune related complications of HCT. Prior to conditioning, neutrophils, erythrocyte progenitors and monocytes were the major contributors of cfDNA in plasma (22.0%, 12.0% and 11.7%, respectively, average cfDNA concentration 208 ± 280 ng/mL plasma). A variety of HCT conditioning regimens have been developed with varying degrees of organ toxicity and myelosuppression. Most patients in our cohort received reduced intensity conditioning therapy (RIC, n=25), whereas two patients received myeloablative conditioning therapy. Comparison of cfDNA tissues-of-origin in plasma before and after conditioning showed a significant drop in blood-derived cfDNA as expected from the function of the conditioning therapy (mean proportion of hematopoietic cell cfDNA decreased from 78% ± 8% to 55% ± 22%, p-value=9.6×10^−5^, **Fig. 2C**). The proportion of blood-derived cfDNA increased to 82% ± 11% at engraftment (p-value = 1.4×10^−5^, **Fig. 2C**). The most notable effect of stem cell infusion and engraftment was a significant increase in the absolute concentration of cfDNA (mean human-derived cfDNA concentration from 190 ng/mL on day of transplant to 1494 ng/mL at engraftment [p-value = 0.020], **Fig. 2D**).

We next evaluated the performance of a cfDNA tissue-of-origin measurement to predict GVHD (**Fig. 2E**). We defined GVHD here as the clinical manifestation of any stage of the disease within the first 6 months post HCT (GVHD+, see Methods). We excluded samples collected after GVHD diagnosis, as these patients received additional GVHD treatment. We found that the concentration of solid-organ specific cfDNA was significantly elevated for patients in the GVHD+ group at month 1, 2 and 3 (p-values of 0.0025, 0.0032, 0.0044, respectively), but not at the two pre-transplant time points (p = 0.71 prior to conditioning, and p = 0.84 prior to hematopoietic cell infusion) (**Fig. 2 E**). Receiver operating characteristic analysis (ROC) of the performance of cfDNA as a predictive marker of GVHD yielded an area under the curve (AUC) of 0.88, 0.95 and 0.96 at engraftment and months 1, 2, and 3, respectively. These results support the notion that cfDNA predicts GVHD occurrence as early as one month after HCT (mean solid organ cfDNA of 872 and 38 ng/mL plasma for GVHD+ and GVHD-, respectively; ROC AUC = 0.88, p-value = 0.0025) (**Fig. 2E**).

To evaluate the ability of this assay to pinpoint the site of GVHD, we quantified the burden of skin-derived cfDNA in the blood of GVHD negative individuals (n=13) and individuals who developed cutaneous GVHD (n=12). We found that plasma samples from individuals with GVHD had a higher burden of skin-derived cfDNA prior to clinical diagnosis of skin GVHD when compared to samples from individuals who did not develop cutaneous GVHD (mean skin cfDNA of 7.1 ng/mL plasma and 2.1 ng/mL plasma, respectively, p-value = 0.047 for samples collected after engraftment and before clinical diagnosis, **supplementary figure 3**). The number of samples from patients diagnosed with hepatic and gastrointestinal GVHD was insufficient to test the performance of the assay to identify GVHD related injury to the liver or gut (n = 3 and n = 5, respectively).

**Figure 3.**
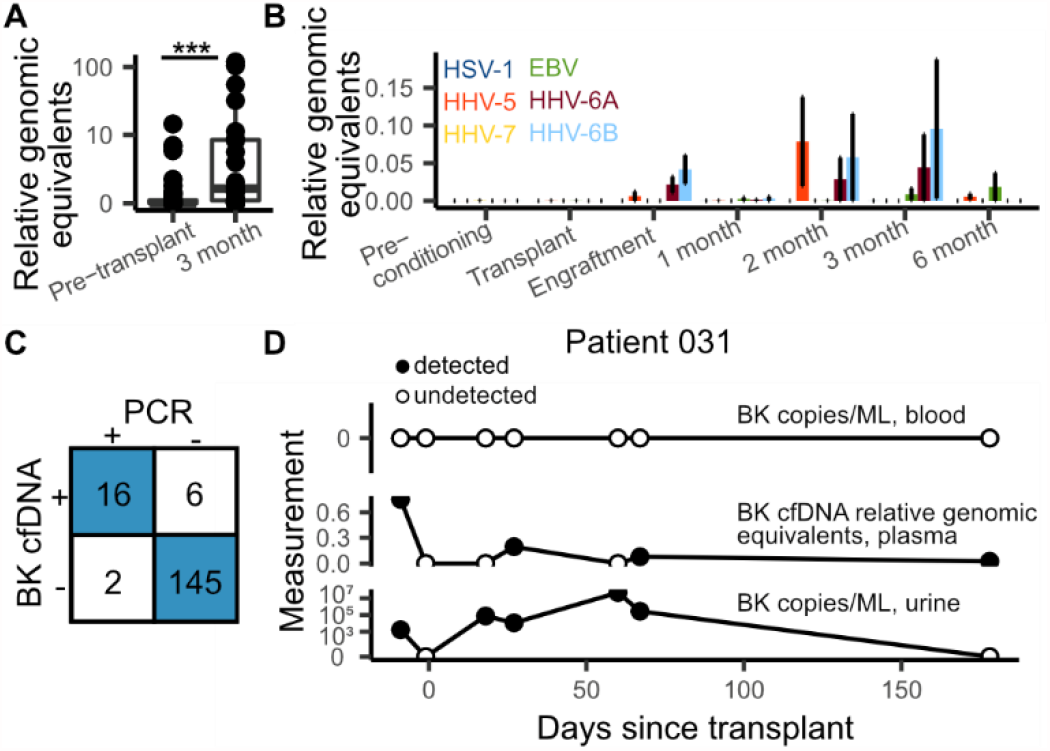
Infectome screening in HCT patients. **A** Relative genomic equivalents of Anelloviruses detected before transplant (pre-conditioning and transplant timepoints) and the 3 month timepoint. **B** Relative genomic equivalents of human herpesviruses by timepoint. Error bars represent standard error of the mean. **C** Concordance between clinically validated BK PCR test (in blood) and BK cfDNA identification. **D** BK abundances in blood (PCR test, top), plasma (cfDNA, middle) and urine (PCR test, bottom) in patient 031.

### Plasma virome screening after HCT

cfDNA from microbes can be detected in the circulation, providing a means to screen for infection via metagenomic cfDNA sequencing^22,33–36^. This may be a particularly powerful approach in the context of HCT, given the high incidence of infectious complications, and the broad range of microorganisms that can cause disease in HCT. To test this concept, we mined all cfDNA data for microbial derived sequences. In a previous study, we found close agreement between the abundance of organisms measured by shotgun sequencing of untreated and bisulfite-treated cfDNA, confirming the possibility to perform metagenomic cfDNA sequencing by WGBS^37^. To identify microbial-derived cfDNA after WGBS, we first identified and removed host related sequences and we aligned the remaining unmapped reads to a set of microbial reference genomes (0.2 ± 0.4% of total reads, Materials and Methods). We implemented a background correction algorithm to remove contributions due to alignment noise and environmental contamination and compared species abundances by the relative abundance of species reads to human reads^35,38^ (relative genomic equivalents [RGE]).

Using this procedure, we found a significant increase after HCT in the burden of cfDNA derived from DNA viruses (DNA sequencing is not sensitive to RNA molecules, average RGE of 1.34 and 26.1, Pre-conditioning and month 3, respectively, p-value = 0.0090). Viruses from the *Anelloviridae* family were the most abundant (463 occurrences of an *Anelloviridae* species). We and others have reported a link between the abundance in plasma of *Anelloviridae* and the degree of immunosuppression in transplantation^39,40^. In line with these observations, the increase in cfDNA derived from DNA viruses was largely due to an increase in the burden of *Anelloviridae* in the first months after HCT (**Fig. 3A**). *Herpesviridae* and *Polyomaviridae* frequently establish latent infection in adults and may reactivate after allogeneic HCT^41^. We identified cfDNA from Human *Herpesviridae* and *Polyomaviridae* in 100 of 170 samples from 26 of 27 patients (**Fig. 3B,C**).

BK Polyomavirus PCR tests are routinely performed in this patient population due to the frequent complications related to BK Polyomavirus. We tested the sensitivity of the cfDNA assay against a clinically validated BK Polyomavirus PCR screening test and found strong concordance (sensitivity = 0.89, specificity = 0.96). For 4/6 discordant readouts, where the cfDNA test detected BK polyomavirus and the PCR test did not, three were from a patient with clinically confirmed reactivation of the virus in the urine (**Fig. 3D**). These findings demonstrate the possibility to sensitively screen for infectious complications after HCT via cfDNA.

### Tumor-specific and donor-specific cell-free DNA inform cancer relapse and loss of engraftment

Many studies have established the utility of circulating tumor-specific cfDNA for early cancer detection and monitoring of minimal residual disease. Here, we assessed the utility of cfDNA profiling of cancer-associated copy number alterations (CNAs) as an approach to detect the presence of leukemia-derived DNA in plasma. At the Dana-Farber Cancer Institute, chromosomal aberrations related to malignant blood disorders are examined using a clinically-validated, targeted, ultra-deep sequencing assay (pre transplant PBMC, n = 20 patients, Rapid Heme Panel, RHP^21^). Using RHP data, we identified six patients with CNAs. We next analyzed all cfDNA WGBS sequence data and found the cfDNA assay was able to detect leukemia-specific CNAs before transplant in two of these patients (**Fig. 4A-C**, patients 003 [mortality] and 031 [no mortality]). Relative copy number changes were used to estimate the fraction of cell-free originating from tumor cells (**Fig. 4B**, see Methods). Last, we found that the genome-wide cfDNA assay enabled detection of CNAs in regions not included in the Rapid Heme Panel (**Fig 4D)**, underlining the importance of a genome-wide approach^42^.

**Figure 4.**
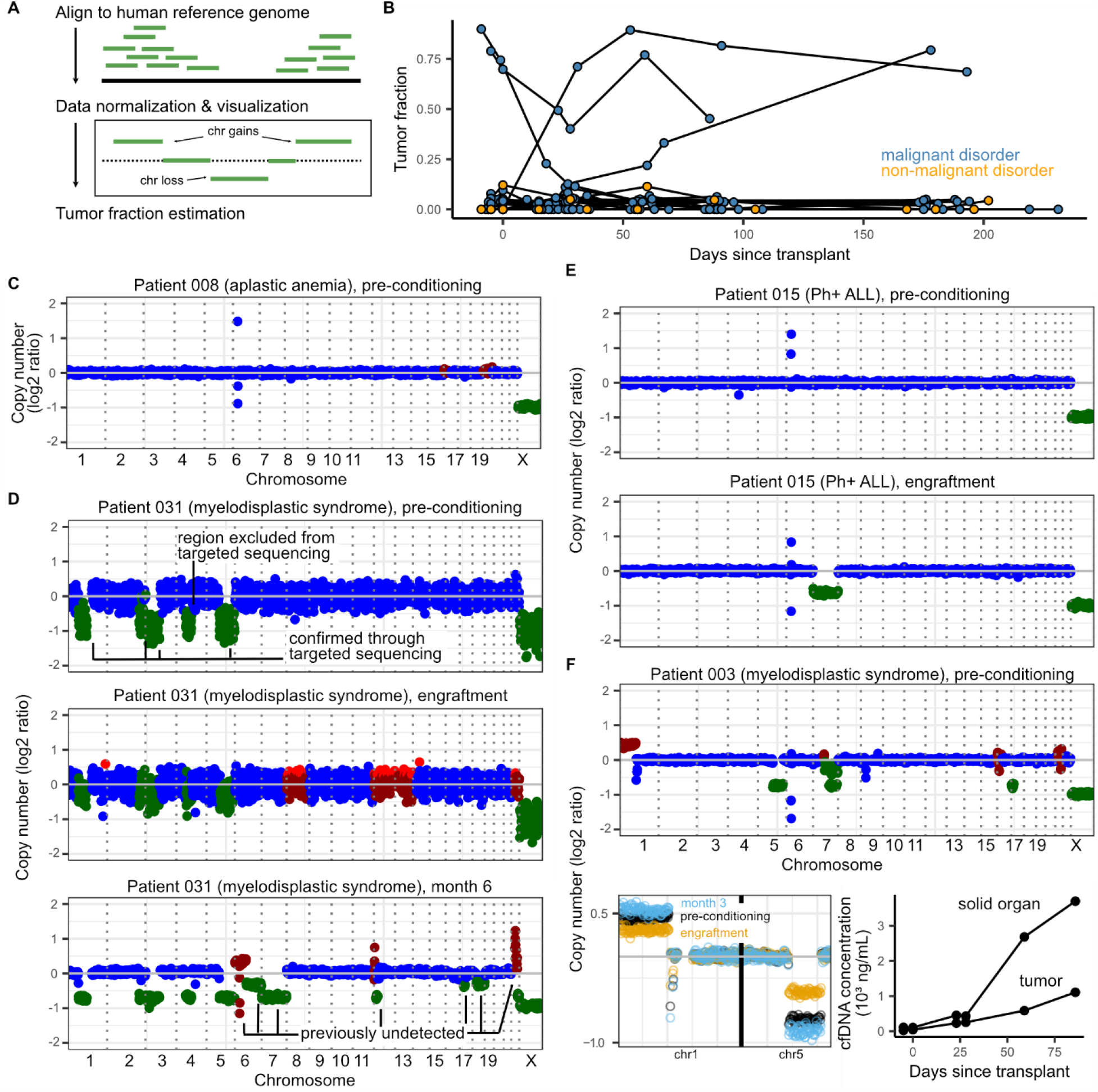
**A** Overview of tumor fraction estimation using copy number alterations. **B** Tumor fractions as measured through ichorCNA at all collected timepoints. Patients without malignant disease and without CNAs (as identified through targeted sequencing) were used to gauge the error in tumor fraction measured by ichorCNA (up to 12%). **C** Example of a copy number alteration profile in a patient with a non-malignant blood disorder (with no alterations expected). The few outliers in the coverage plot for patient 008 are likely due to errors in sequence mapping. Genome-wide plots in C-F (top only in F) are colored by ichorCNA’s identification of a given region as neutral (blue), gained (red) or lost (green). **D-F** Copy number alteration profiles of three patients with measurable copy number alteration-based tumor fractions. **D** Patient 015 was found to have loss of chromosome 7 at the time of engraftment and in all subsequent samples. **E** Patient 031, over the course of their treatment, developed additional, clinically undetected structural variants. **F** Patient 003 (deceased on day 91) had detectable tumor fraction and clinical evidence of GVHD. Solid-organ derived cell-free DNA was higher than the tumor load (line plot, right-hand side). Top: genome-wide coverage plot. Bottom left: copy number profiles on chromosomes 1 and 5 show a decrease in copy number changes at engraftment (yellow) and subsequent increase at month 3 (blue), when compared to the pre-conditioning timepoint (black). Bottom right: Tumor and solid organ derived cfDNA concentration at all available timepoints for patient 003. Patients 003, 008, 015 and 031 were all male-male donor-recipient pairs.

We highlight three cases that exemplify the utility of continuous patient monitoring. First, cfDNA monitoring for patient 031 detected new CNAs after HCT, suggesting the expansion of a subclonal tumor population over the course of treatment (**Fig. 4D, supplementary figure 4)**. In this patient, we estimated tumor fractions of 90% at pre-conditioning, 23% at engraftment, and 79% at month 6. Bone marrow biopsies performed 8.5 months after transplant (outside the timeframe of the current study) revealed hypocellular marrow consistent with acute myeloid leukemia. Second, profiling of patient 015 diagnosed with Philadelphia chromosome positive ALL (Ph+ALL) revealed the presence of monosomy 7 at engraftment and all subsequent time points (**Fig. 4E, supplementary figure 5**). Clinical chimerism testing based on short-tandem repeat PCR amplification for this same patient showed full engraftment of donor cells and bone marrow examination showed no evidence of leukemia relapse. Cytogenic analysis performed after transplant confirmed monosomy 7 in donor cells for this patient, highlighting the utility of an untargeted sequencing assay to identify rare transplantation events. Last, for patient 003 (**Fig. 4F, supplementary figure 6)** who was diagnosed with severe GVHD (cutaneous stage 4, overall grade IV; unresolved; mortality day 91), cfDNA tissue-of-origin profiling revealed an increase in solid-organ derived cfDNA in addition to increasing tumor-derived cfDNA load (**Fig. 4F**), potentially pointing towards a joint graft-versus-host disease and relapse.

### Plasma donor-derived cfDNA as a marker of mixed chimerism

Measurements of donor-recipient chimerism are a routine part of clinical monitoring and can inform cancer relapse and loss the donor stem cell graft^43,44^. These measurements are performed on isolated hematopoietic cell populations and do not account for the turnover rate of cells, which are often higher in leukemic cells than in normal cells^45^. Therefore, it has been proposed that chimerism analysis of cfDNA may offer complimentary information to traditional cell-based chimerism analysis^17,46^. Here, we show the feasibility of measuring donor-derived cfDNA (dd-cfDNA) by leveraging the relative abundance of X and Y chromosomes in sex-mismatched recipient pairs (samples with depth of sequencing > 0.1x, **Fig. 5A**). We analyzed samples collected prior to transplantation to assess the error rate of this measurement (mean donor fraction 0.0% ± 4.6%). We found that the donor fraction is highest at engraftment (86%% ± 13%) and remains constant in the absence of complications (**Fig. 5B, supplementary figure 7**). We highlight two examples from patients who experienced HCT complications (**Fig. 5C**). In Patient 002 we observed a gradual decrease in dd-cfDNA after engraftment, with a sharp drop on day 183. This patient developed disease relapse on day 152 and the gradual decreased in dd-cfDNA preceded relapse. Similarly, in patient 006, we observed a steady drop in dd-cfDNA after engraftment, prior to disease relapse on day 85. The fraction of dd-cfDNA for this patient subsequently increased at month six, before the patient entered remission (day 218). Taken together, these data suggest that dd-cfDNA is an informative biomarker for HCT monitoring and can be used in conjunction with other cfDNA features to inform levels of donor cell engraftment and quantification of residual recipient cells and relapse.

**Figure 5.**
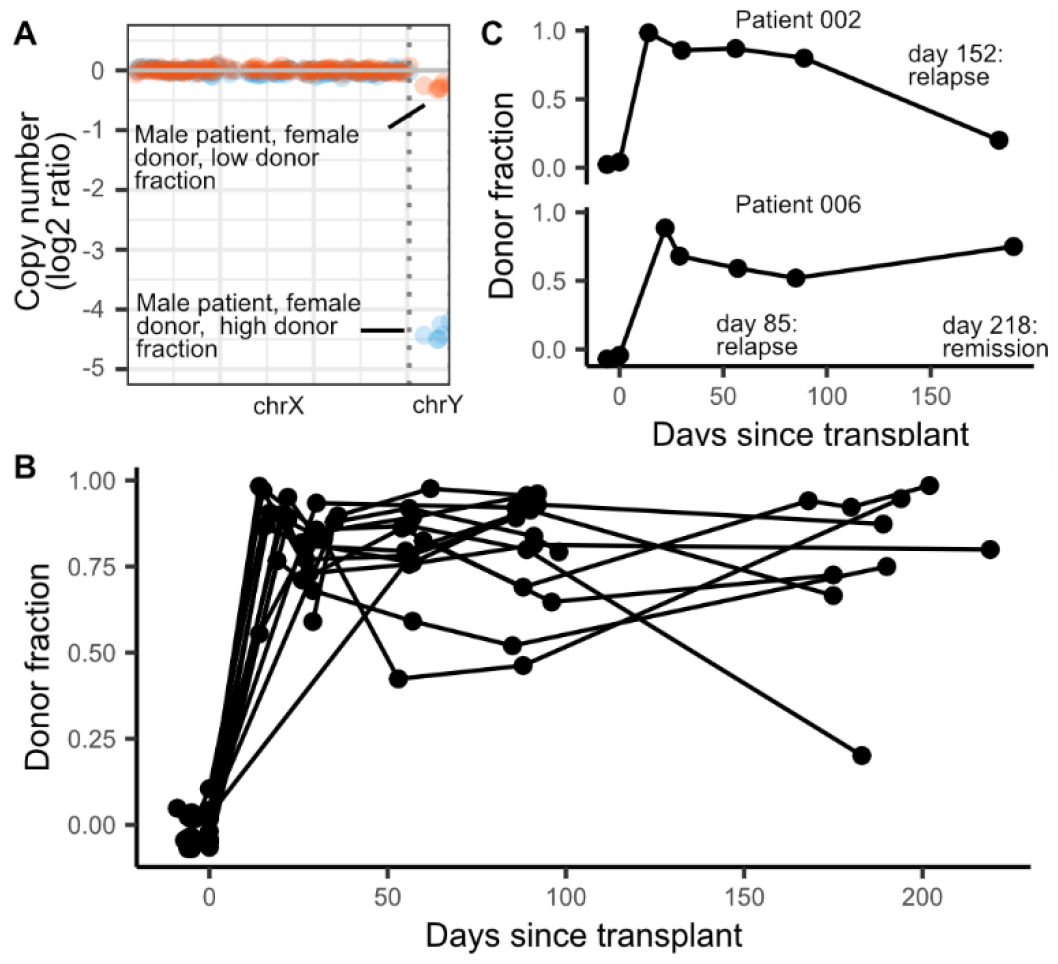
Donor fractions and days post-transplant in sex-mismatched patients. **A** The donor fraction is measured from the relative coverage of sex chromosomes (see Methods). **B** Donor fraction in all sex-mismatched patients. **C** Donor fraction in two patients who experienced disease relapse.

## DISCUSSION

In this work, we have introduced a cfDNA assay with the potential to simultaneously screen for the most important complications that arise after allogeneic HCT. This work was inspired by recent studies that have shown that cfDNA is an analyte with utility in *i)* monitoring rejection after solid-organ transplantation^47–50^; *ii)* screening for infection and viral reactivation^33,36,37^ and *iii)* early detection of cancer or relapse of disease^42,51,52^.

Numerous studies have demonstrated that donor derived cfDNA in the blood of solid-organ transplant recipients is a quantitative marker of solid-organ transplant injury^48,49^ and a variety of commercial cfDNA assays are already in use^47,53–56^. We reasoned that cfDNA may also inform tissue injury due to GVHD after HCT. To quantify cfDNA derived from any tissue, we implemented bisulfite sequencing of cfDNA, to profile cytosine methylation marks which are comprised within cfDNA and are cell, tissue and organ type specific. We found that the burden of cfDNA from solid organs is predictive of the onset of GVHD as early as one month after HCT. Protein biomarkers have previously been investigated for diagnosis and prediction of GVHD^57,58^. ST2 and REG3α, which both derive from the gastrointestinal tract, are two such biomarkers with the strongest predictive power. The cfDNA assay presented here has inherent advantages over these protein biomarker technologies. First, cfDNA may provide a generalizable approach to measure injury to any tissue, whereas protein injury markers may not be available for all cell and tissue types. Second, because the concentration of tissue-specific DNA can be directly related to the degree of cellular injury^37,59–61^, cfDNA may offer a measure of injury that can be trended over time.

Whole genome sequencing is not only responsive to human host-derived cfDNA, but also to microbial cfDNA in the blood circulation. Several recent studies have demonstrated the value of metagenomic cfDNA sequencing to screen for infection in a variety of clinical settings, including urinary tract infection^33,37^, sepsis^36^, and invasive fungal disease^62^. In HCT, metagenomic cfDNA sequencing has been used to identify pathogens in blood before clinical onset of bloodstream infections^63^. Here, we explored the potential to identify viral derived cfDNA in plasma of HCT recipients using whole genome bisulfite sequencing. This approach revealed the frequent presence of cfDNA from anelloviruses, cytomegalovirus, herpesvirus 6, Epstein-Barr virus and polyomavirus in the blood of HCT recipients. Anelloviruses were common in this cohort, and, while rarely pathogenic, can be used as a surrogate for the degree of immunosuppression in transplant patients^39,40,64^. We demonstrate sensitive detection of BK virus cfDNA for patients that were BK virus positive in blood, and for a patient that tested negative for BK virus in the blood, but tested positive for BK virus in the urine, which may indicate that the cfDNA assay reported here has a higher sensitivity than clinical PCR assays. The assay reported here has the potential to simultaneously inform about GVHD, from the tissues-of-origin of host cfDNA, and infection, from metagenomic analysis of microbial cfDNA. Compared to conventional metagenomic sequencing, this assay requires one additional experimental step to bisulfite convert cfDNA, which can be completed within approximately 2 hours and is compatible with multiple existing next-generation sequencing workflows.

Circulating tumor DNA has been shown to be a highly sensitive molecule for the detection of minimal residual disease^42,65^. The identification of solid-tumor derived circulating nucleic acids relies on the identification of single-nucleotide polymorphisms or copy number alterations^51,66^, or detection of changes in DNA fragmentation patterns^52,67,68^,. In this work, we focused on structural variants of malignant disease to detect tumor-specific cfDNA and found evidence of subclonal expansion, newly acquired mutations, and simultaneous occurrence of GVHD and cancer relapse. Future studies where whole-genome sequencing is performed on the primary tumor cells may uncover tumor-associated SNPs and be used in conjunction with CNA analysis to improve detection of circulating tumor DNA in malignant blood disease^42^.

Donor-derived cells and, recently, dd-cfDNA have been explored as markers for GVHD, loss of graft and recurrence of disease^19,46^. We observed increased amounts of dd-cfDNA at engraftment, and these levels remained elevated in the absence of HCT complications. For patients that suffered relapse of disease, we observed a decrease in the burden of dd-cfDNA, potentially due to suppression of normal marrow cells by leukemic cells or increases in recipient tissue damage^19,69^. Studies by Duque-Afonso *et al* and Sharon *et al* reported elevated amounts of transplant recipient cfDNA in cases of GVHD, suggesting patient tissue contributions to the cell-free DNA mixture^19,46^. Interestingly, Duque-Afonso *et al* also observed increased recipient cfDNA at the time of relapse and progressive disease, suggesting that donor-derived (or recipient-derived) cfDNA alone may not be sufficient in distinguishing different important complications of HCT, supporting the need for an assay that is informative of the tissues-of-origin of cfDNA.

This is a proof-of-principle study with several limitations that can be addressed in future work. First, the scope of the current study with 27 patients was not powered to detect any association of cfDNA with acute GVHD involving organs other than skin (liver, gut). Our results suggest that cfDNA tissue-of-origin profiling is predictive of acute GVHD, but larger studies will be needed to extend the current observations to other sites of organ damage, and to assess its utility in detecting and diagnosing chronic GVHD. In addition, larger studies, including patient populations with diverse HCT complications, are necessary to resolve the origins of cell-free DNA in cases of relapse of disease. Despite these potential limitations, we have shown here that cell-free DNA is a versatile analyte to monitor HCT patients, and our data highlights the importance of comprehensive monitoring all origins of cell-free DNA to assess the most severe complications of HCT.

## MATERIALS AND METHODS

### Study cohort

We performed a nested case-control study within a prospective cohort of adult patients undergoing allogeneic HCT at Dana-Farber Cancer Institute. Patients were followed for 6 months after HCT. Patients were selected for this study on a rolling basis, and were placed in the GVHD case or control groups based on clinical manifestation of the disease within the first 6 months after HCT. The study was approved by the Dana-Farber/Harvard Cancer Center’s Office of Human Research Studies. All patients provided written informed consent.

For this study, we used 170 blood samples collected from 27 allogeneic HCT recipients from August 2018- to August 2019. Baseline patient characteristics were recorded. Covariates of interest included HLA matching, donor relatedness and donor-recipient sex mismatch (**supplementary table 1**). Date of onset of GVHD, as well as GVHD prophylaxis and treatment regimens were documented. GVHD was diagnosed clinically and pathologically. GVHD severity was graded according to the Glucksberg criteria<u>^43^</u>. Other clinical events of interest included the development of bloodstream infections, BK polyomavirus disease, and clinical disease from other DNA viruses.

### Time points

Standard time points for plasma collection were determined prior to patient recruitment and included pre-conditioning (the day of their first conditioning dose, prior to receiving treatment), transplantation (the day of transplantation, prior to transfusion), engraftment (detailed below) and months 1, 2, 3 and 6 after transplant. In the event of BK-related symptoms, disease or reactivation, additional time points were collected. In the case of two time points overlapping, the sample was preferentially labeled as engraftment, month 1, month 3, or month 6 (in that order).

### Engraftment

Neutrophil engraftment was considered when blood samples contained an absolute neutrophil count greater or equal than 500 cell per microliter of blood on two separate measurements.

### Relapse

Disease relapse was defined through standard criteria for each underlying disease.

### Mixed chimerism

Mixed chimerism is broadly defined as 5-95% T cells of donor origin^43^. Here, we used a criteria of <75% T cells of donor origin to characterize mixed chimerism. Only timepoints obtained after engraftment were considered.

### BK polyomavirus disease identification

Patients were identified as BK virus disease positive when they presented BK-related urinary symptoms that correlated with positive BK qPCR test in either urine or blood (>10^5^ copies/ mL in urine, >0 copies/mL in blood; Viracor BK qPCR test, reference #2500) and did not have evidence of any other cause of genitourinary pathology at the time of symptom onset.

### Blood sample collection and plasma extraction

Blood samples were collected through standard venipuncture in EDTA tubes (Becton Dickinson (BD), reference #366643) on admission, before the beginning of the conditioning chemotherapy; on the day of HCT after the completion of the conditioning chemotherapy, at engraftment (usually 14 to 21 days after HCT), and at months 1, 2, 3 and 6 post-HCT. Plasma was extracted through blood centrifugation (2000rpm for 10 minutes using a Beckman Coulter Allegra 6R centrifuge) and stored in 0.5-2mL aliquots at –80 °C. Plasma samples were shipped from DFCI to Cornell University on dry ice.

### Nucleic acid control preparation

Synthetic oligos were prepared (IDT, **supplementary table 2**), mixed in equal proportions, and diluted at approximately 150 ng/ul. At the time of cfDNA extraction, 8ul of control was added to 1992µL of 1xPBS and processed as a sample in all downstream experiments.

### Cell-free DNA extraction

cfDNA was extracted according to manufacturer recommendations (Qiagen Circulating Nucleic Acid Kit, reference #55114, elution volume 45µl). Eluted DNA was quantified using a Qubit 3.0 Fluorometer (using 2µL of eluted DNA). Measured cfDNA concentration was obtained using the following formula:

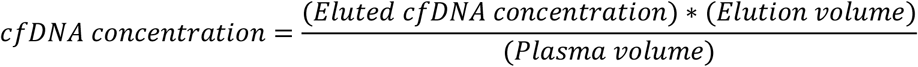

### Whole-genome bisulfite sequencing

cfDNA and nucleic acid controls were bisulfite treated according to manufacturer recommendations (Zymo Methylation Lightning Kit, reference #D5030). Sequencing libraries were prepared using a previously described single-stranded library preparation protocol. Libraries were quality-controlled through DNA fragment analysis (Agilent Fragment analyzer) and sequenced on an Illumina NextSeq550 machine using 2×75bp reads. Nucleic acid controls were sequenced a ∼1% of the total sequencing lane.

### Human genome alignment

Adapter sequences were trimmed using BBTools^70^. The Bismark alignment tool^71^ was used to align reads to the human genome (version hg19), remove PCR duplicates and calculate methylation densities.

### Reference tissue methylation profiles and tissue of origin measurement

Reference tissue methylomes were obtained from publicly available databases^28–32^ (**supplementary dataset 1**). Genomic coordinates from different sources were normalized and converted to a standard 4 column bed file (columns: chromosome, start, end, methylation fraction) using hg19 assembly coordinates. Methylation profiles were grouped by tissue-type and differentially methylated regions were found using Metilene^72^. Tissues and cell-types of origin were determined using quadratic programming as previously described^37^.

### Donor fraction

Donor fractions were calculated by measuring the relative coverage of X and Y chromosomes in sex-mismatched donor-recipient pairs. Coverage was summed across binned, 500 base pair windows and adjusted for mappability and GC content using HMMcopy^33,73^.

### Tumor fraction

ichorCNA^66^ (version 2.0) was used to detect copy number alterations and estimate tumor fraction in patients with cancer. A window size of 1MB along with a ploidy of (2,3) and a wide range of non-tumor restart fractions were used to calculate coverage on autosomal chromosomes. Coverage was normalized using a panel of normals generated from the plasma of 5 healthy donors (IRB XYZ). The plasma used for the panel of normals was processed using the same workflow as described above to account for experimental and sequencing artifacts. The normalized coverage profile for each sample was then used to detect copy number alterations and estimate tumor fraction.

### Metagenomic alignment and quantification of microbial cfDNA

After WGBS, reads were adapter-trimmed using BBTools^70^, and short reads are merged with FLASH^74^. Sequences were aligned to a C-to-T converted genome using Bismark^71^. Unmapped reads were BLASTed^75^ using hs-blastn^76^ to a list of C-to-T converted microbial reference genomes. A relative abundance of all detected organisms was determined using GRAMMy^77^, and relative genomic abundances are measured as previously described^35^. Microbial cfDNA fraction was calculated by dividing the unique number of reads mapping to microbial species (after adjusting for the length of each microbial genome in the reference set) to the total number of adapter-trimmed reads. Human fraction is estimated as 1 - microbial fraction. Microbial species were then filtered for environmental contamination and alignment noise using previously described methods^38^.). **cfDNA concentration**. cfDNA concentration of a specific tissue or microbe is calculated as follows:

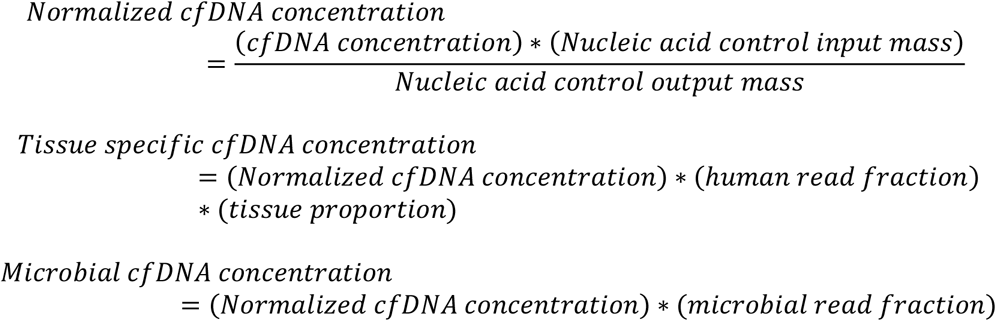

### Depth of coverage

The depth of sequencing was measured by summing the depth of coverage for each mapped base pair on the human genome after duplicate removal, and dividing by the total length of the human genome (hg19, without unknown bases).

### Bisulfite conversion efficiency

We estimated bisulfite conversion efficiency by quantifying the rate of C[A/T/C] methylation in human-aligned reads (using MethPipe^78^), which are rarely methylated in mammalian genomes.

### Statistical analysis

Statistical analysis was performed in R (version 3.5). All tests were performed using a two-sided Wilcoxon test.

## Data availability

The sequencing data generated for this study will be available in the database of Genotypes and Phenotypes (dbGaP). All code used to generate figures and analyze primary data will be made available on GitHub.

## AUTHOR CONTRIBUTIONS

A.P.C., M.P.C., F.M.M., J.R. and I.D.V. designed the study. A.P.C. and J.S.L. performed experiments. M.P.C., F.M.M., J.R., K.C., K.M.T., J.L.O. and E.S. consented patients and acquired clinical data. A.P.C., C.J.L., S.S., M.P.C., P.B., I.D.V., F.M.M. and J.R. analyzed data. A.P.C., M.P.C., I.D.V., F.M.M. and J.R. wrote the manuscript. All authors reviewed and approved the manuscript.

## ACKNOWLEDGEMENTS

We would like to express our gratitude and deepest respect for our co-author and colleague, Dr. Francisco M. Marty who passed away in April 2021. He was an Associate Professor in the Department of Medicine at the Brigham and Women’s Hospital, Dana-Farber Cancer Institute and Harvard Medical School. Francisco had an unrivaled passion for patient care and clinical research focusing on the treatment of opportunistic infections in immunocompromised hosts. He was honored with numerous awards and recognitions for his contributions to teaching, clinical care, and rigorous scientific research during the course of his illustrious career. He was a great friend and mentor to many trainees and collaborators across the world. He will be dearly missed.

We would like to thank the Cornell Genomics Center for help with sequencing assays, the Cornell Bioinformatics facility for computational assistance, the Pasquarello Tissue bank at the Dana-Farber Cancer Institute for sample processing and cryopreservation and members of the De Vlaminck Lab for helpful discussions. We thank Francoise Vermeulen of the Cornell Statistical Consulting Unit for helpful discussion. This work was supported by R01AI146165 (to I.D.V, J.R, M.P.C. and F.M.), R21AI133331 (to I.D.V.), R21AI124237 (to I.D.V.), DP2AI138242 (to I.D.V.), a National Sciences and Engineering Research Council of Canada fellowship PGS-D3 (to A.P.C)

